# A CRISPR screen reveals a WNT7B-FZD5 signaling circuit as a therapeutic opportunity in pancreatic cancer

**DOI:** 10.1101/041996

**Authors:** Zachary Steinhart, Traver Hart, Megha Chandrashekhar, Zvezdan Pavlovic, Mélanie Robitaille, Xiaowei Wang, Jarrett Adams, James Pan, Sachdev Sidhu, Jason Moffat, Stéphane Angers

**Author notes:** Correspondence to (S.A), (J.M), (S.S.).

## Abstract

CRISPR-Cas9 genome editing enables high-resolution detection of genetic vulnerabilities of cancer cells. We conducted a genome-wide CRISPR-Cas9 screen in *RNF43* mutant pancreatic ductal adenocarcinoma (PDAC) cells, which rely on Wnt signaling for proliferation, and discovered a unique requirement for a WNT7B-FZD5 signaling circuit. Our results highlight an underappreciated level of functional specificity at the ligand-receptor level. We derived a panel of recombinant antibodies that reports the expression of nine out of ten human Frizzled receptors and confirm that WNT7B-FZD5 functional specificity cannot be explained by protein expression patterns. We developed two human antibodies that target FZD5 and robustly inhibited the growth of *RNF43* mutant PDAC cells grown *in vitro* and as xenografts, providing strong orthogonal support for the functional specificity observed genetically. Proliferation of a patient-derived PDAC cell line harboring a *RNF43* variant previously associated with PDAC was also selectively inhibited by the FZD5 antibodies, further demonstrating their use as a potential targeted therapy.

## Introduction

Wnt signaling pathways regulate proliferation and differentiation of stem and progenitor cells during embryonic development and adult tissue homeostasis^1,2^. Deregulation of Wnt-β-catenin signaling has been linked to tumor initiation and maintenance in several human malignancies^3^. In most tumors Wnt-β-catenin signaling is activated as a result of inactivating mutations in the negative regulators APC and Axin or activating mutations in β-catenin^4^. In these cases the signaling pathway is activated distally from the Frizzled-LRP5/6 Wnt receptor complex at the cell surface and therefore developing drugs against the Wnt pathway has proven challenging^5^.

Recent sequencing efforts of gastric, ovarian and pancreatic neoplasias, in addition to colorectal adenocarcinoma and endometrial carcinoma, has identified recurrent non-synonymous *RNF43* mutations^6-11^ that appear to be mutually exclusive with APC and β-catenin mutations. *RNF43* and its homologue *ZNRF3* encode transmembrane E3 ubiquitin ligases targeting Frizzled receptors, whose loss-of-function mutations lead to high Frizzled expression at the cell surface and sensitize tumor cells to Wnt-dependent growth^11,12^. Epithelial organoids derived from tumors isolated from *RNF43*^-/-^;*ZNRF3*^-/-^ mice grow in the absence of R-Spondins, highlighting their hypersensitivity to Wnt. Importantly, the growth of the same organoids is blocked by inhibition of Porcupine (PORCN), an O-acyltransferase required for the maturation and activity of Wnt proteins, indicating a requirement for Wnt ligands^13^. The findings showing that treatment of *RNF43*^-/-^;*ZNRF3*^-/-^ mutant mice with a PORCN inhibitor represses the growth of intestinal tumors while leaving adjacent normal crypt intact, suggest that a therapeutic window exists to block overactive Wnt pathway activity upstream of β-catenin in *RNF43* or *ZNRF3* mutant cancers^13^.

The CRISPR-Cas9 system has enabled the simple and efficient genome editing of cultured cells and living organisms and has unlocked the power of genetic screens in human cells^14^. We and others have developed lentiviral-based pooled gRNA libraries to perform genetic screens to identify essential genes in human cells^15-18^. This work uncovered ~2,000 essential genes in each cell line, including a common core of ~1,600 essential genes^15,17^. Strikingly, ~400 genotype-specific or context-dependent fitness genes were also identified for each of the cell lines, exposing vulnerabilities that may be tractable therapeutic targets^15,17^. Because of the urgent need for new therapeutic strategies for pancreatic cancer, we applied genome-scale pooled CRISPR screening technology to identify vulnerabilities in *RNF43*-mutant pancreatic adenocarcinomas.

## Results

To identify context-dependent fitness genes in *RNF43*-mutant pancreatic cancer cells, we used the HPAF-II PDAC cell line that was previously shown to be exquisitely sensitive to PORCN inhibition^19^. We carried out a genetic screen using the TKO gRNA library and monitored evolving cell populations over ~20 doublings by deep sequencing of gRNAs (Table S1). gRNA abundance over multiple time points was assessed using gold-standard sets of essential and nonessential genes^20^. The fold-change distribution of gRNAs targeting essential genes was significantly shifted relative to those targeting nonessential genes, and this shift increased with time indicating that the screen functioned as designed (Fig. 1a). We then used the BAGEL algorithm^15,21^ to calculate a log Bayes Factor (BF) for each gene, which is a measure of the confidence that knockout of a specific gene causes a decrease in fitness (high BF indicates increased confidence that the knockout of the gene results in a decrease in fitness) (Fig. S1 and Table S2). Comparing the HPAF-II screen to TKO fitness screens carried out in five diverse human cell lines^15^ indicated the fitness gene profile of HPAF-II cells was most similar to DLD-1 and HCT116 colorectal cancer cells (Fig. S2a-b), which may reflect the common endodermal origin of these cell lines. A total of 2,174 fitness genes were identified in HPAF-II cells (FDR<5%), including 1,315 of 1,580 (83%) previously identified fitness genes^15^.

**Fig. 1.**
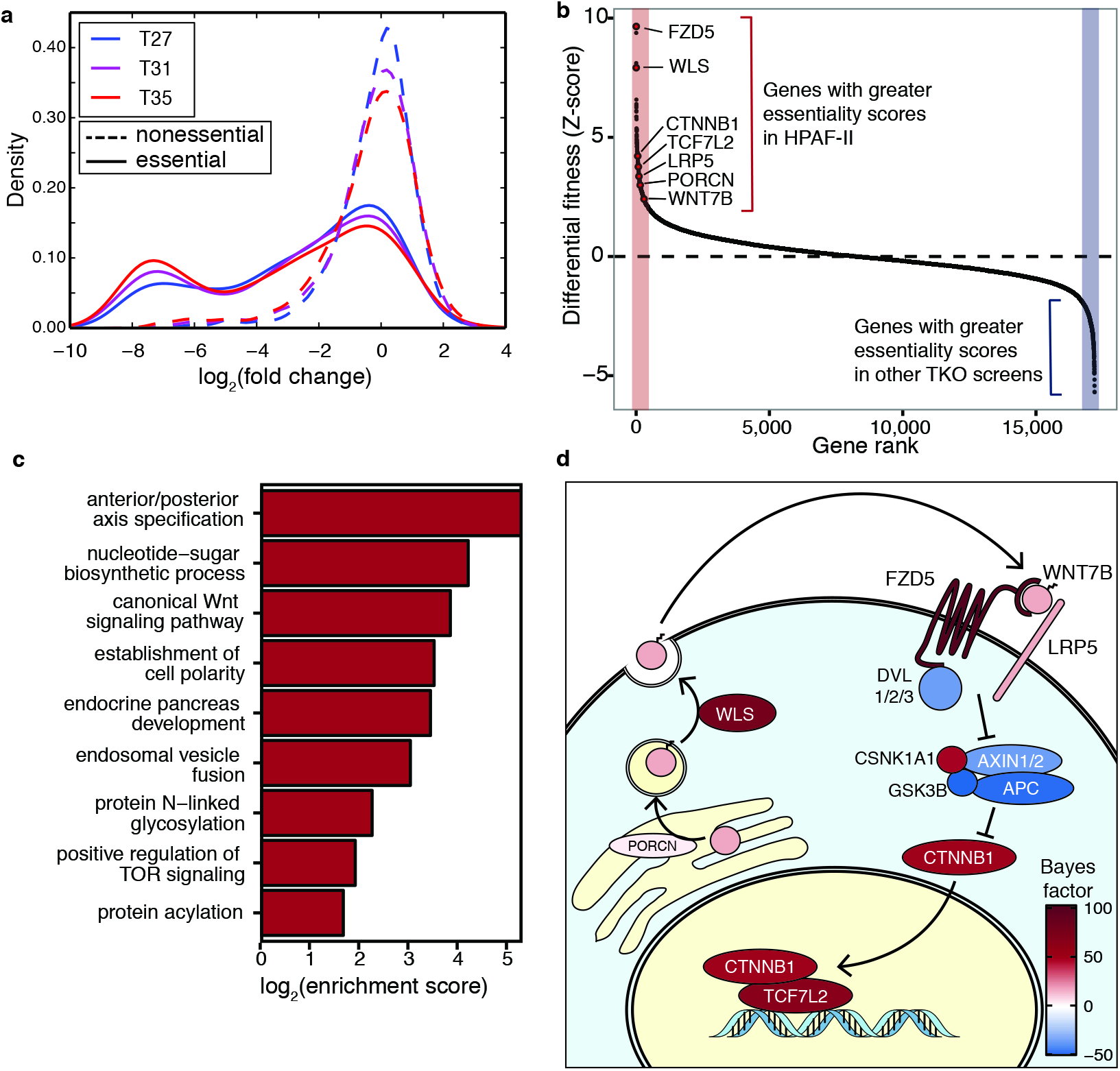
A genome-wide CRISPR/Cas9 screen identifies genetic vulnerabilities of *RNF43* mutant PDAC cells. **(A)** Fold change distributions of gRNA targeting essential genes (solid lines) or nonessential genes (dashed lines) at the indicated time points after infection. **(B)** Ranked differential fitness score reveals context dependent lethality in HPAF-II. BF from HPAF-II were compared to mean BF from HeLa, HCT116, DLD-1, RPE-1 and GBM and converted to a Z-score. **(C)** Selected GO biological process terms enriched among genes ranked in (B). **(D)** Genes essential for Wnt signal transduction in HPAF-II. In this context, autocrine *WNT7B* binds to the *FZD5*/*LRP5* receptor complex, which through the action of *DVL1/2/3* (all three nonessential), inhibits the destruction complex, composed of *AXIN1/2*, *APC*, *GSK3B* and *CSNK1A1*. This allows for relief on inhibition on β-catenin (*CTNNB1*), which can now translocate to the nucleus and activate the *TCF7L2* transcription factor. Each protein is shaded according to its BF or average BF when multiple isoforms are present (e.g. *DVL1/2/3* and *AXIN1/2*)

We next examined the context-dependent fitness genes that were specific to HPAF-II cells compared to other cell lines screened with the TKO library. For each gene, we calculated the difference between Bayes Factor (BF) scores in HPAF-II cells and the average BF scores across the 5 previously reported screens, and converted that difference to a Z-score (Table S3). Examination of the top differential fitness genes readily highlighted the known addiction of HPAF-II cells to Wnt-β-catenin signaling, since we observed several genes previously described as positive regulators of this pathway having Z-scores of ≥2 (*FZD5, WLS, CTNNB1* (β-catenin), *TCF7L2, LRP5, PORCN, WNT7B*) (Fig. 1b). We then analyzed the ranked list of differential essentiality scores for gene ontology (GO) term enrichment and found multiple biological process terms for HPAF-II context-specific essential genes that were related to Wnt biology including “anterior-posterior axis specification”, “canonical Wnt signaling pathway”, and “establishment of cell polarity” (Fig. 1c, Fig. S3). Moreover, HPAF-II context essentials are enriched for the biological processes involved in the biogenesis and activity of Wnt signaling components. These processes include protein N-linked glycosylation, nucleotide-sugar biosynthesis (which produces substrates for glycosyltransferases), protein acylation, and endosomal vesicle fusion.

Core negative regulators of the Wnt-β-catenin pathway were found amongst the lowest BFs including *APC*, *GSK3B*, and *ZNRF3*, suggesting that knockout of these genes may provide a proliferation advantage to HPAF-II cells (Fig. S1). Topping the list in the context-dependent fitness analysis was the Wnt receptor *FZD5* (Z-score >10), which was the only one of ten Frizzled homologs essential for growth of these cells. Thus, while we identify many supporting processes, this data has revealed a surprisingly confined set of genes required for transduction of Wnt-β-catenin signaling in HPAF-II PDAC cells (Fig. 1d).

To validate the screen results we first infected HPAF-II cells with lentivirus coding for various gRNAs, selected transduced cells for 48 hours and plated cells in clonogenic growth assays. Knockout of *FZD5* using two independent gRNAs led to robust growth inhibition, comparable to treatment with a *CTNNB1* gRNA or the PORCN inhibitor LGK974 (Fig. 2a). In contrast, cells transduced with a control *LACZ* gRNA or two validated and unique gRNAs for each of *FZD4*, *FZD7* or *FZD8* exhibited normal growth (Fig. 2a-b and Table S4). We next tested whether FZD5 was also required specifically for the growth of other *RNF43* mutant PDAC cell lines and found that *FZD5* gRNAs, but not *FZD7* gRNAs, inhibited the growth of PaTu-8988S and AsPC-1 cells to levels similar to cells treated with LGK974 or transduced with the *CTNNB1* gRNA (Fig. 2c and Fig. S4). In contrast, the growth of PANC-1 and BxPC-3 cell lines, which are insensitive to LGK974^19^, was not inhibited by gRNAs targeting *FZD5*, *FZD7* or *CTNNB1* (Fig. 2c and Fig. S4). Consistent with these results and supporting a key role for FZD5 in transducing autocrine Wnt-β-catenin signaling in *RNF43* mutant cells, knockout of *FZD5* led to marked inhibition of the Wnt target genes *AXIN2* and *NKD1* whereas minimal or no change was observed in cells transduced with *FZD7* gRNAs (Fig. 2d-e). Furthermore, knocking out *FZD5* or *CTNNB1*, or treatment of cells with LGK974, led to increased expression of the differentiation marker *MUC5AC ^19^*, whereas no change was observed in *RNF43* mutant PDAC cells knocked out for *FZD7* (Fig. 2f). Notably, *WNT7B* was the only Wnt gene out of 19 with a differential fitness Z-score >2 in the genome-wide CRISPR screen (Fig. 1b). Taken together, these results suggest that autocrine WNT7B-FZD5 signaling is responsible for the bulk of β-catenin signaling in *RNF43* mutant PDAC cells. Consistent with this prediction, transduction of a *WNT7B* gRNA strongly inhibited proliferation of HPAF-II cells (Fig. 2g). We conclude that a WNT7B-FZD5 signaling circuit is specifically required for Wnt-β-catenin signaling and that blocking FZD5 or WNT7B is sufficient to inhibit proliferation of *RNF43*-mutant PDAC cells.

**Fig. 2.**
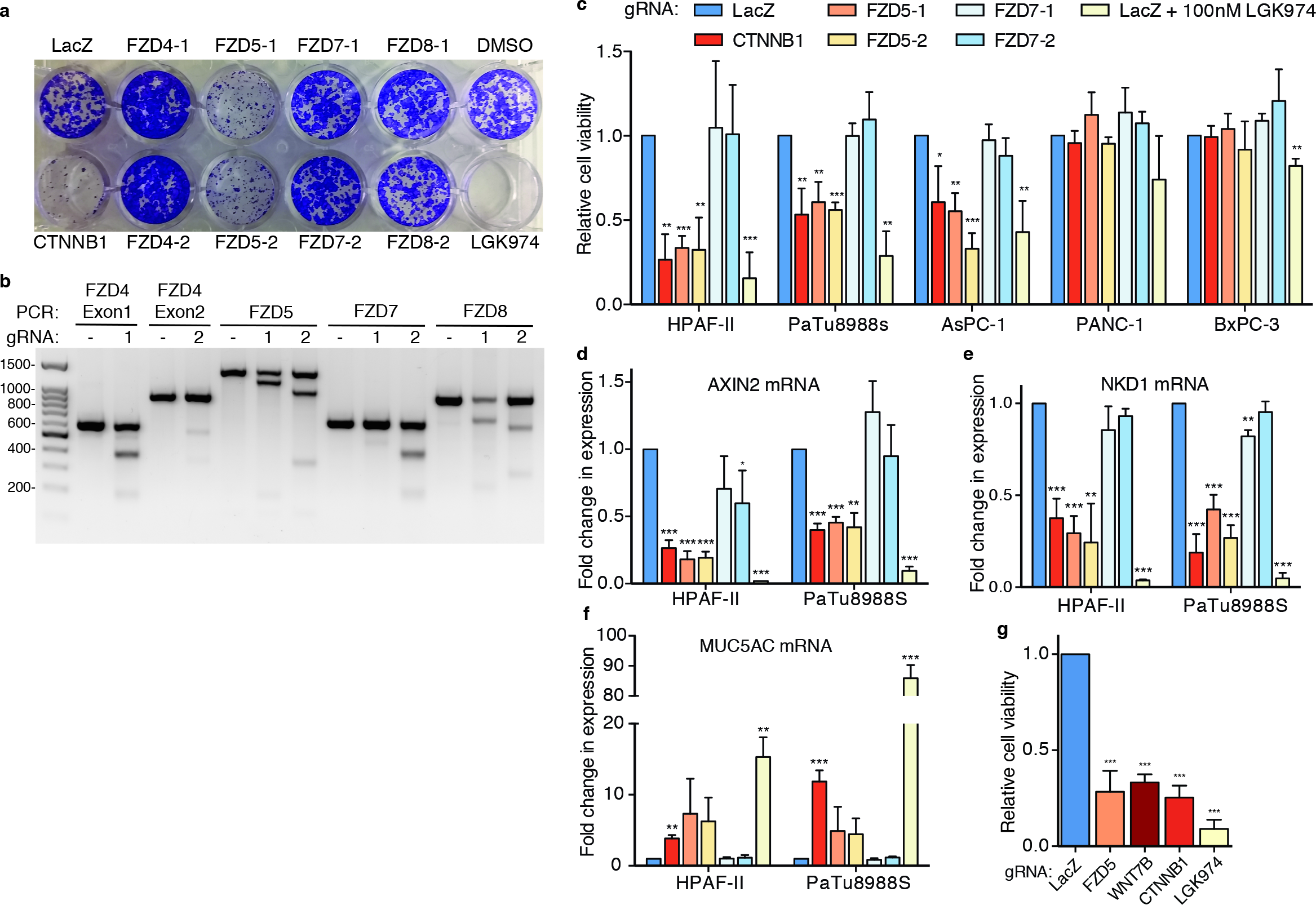
*FZD5* knockout inhibits proliferation of *RNF43* mutant PDAC cell lines and Wnt target genes activation. **(A)** Proliferation assay in HPAF-II cells stably expressing Cas9 and transduced with lentivirus delivering indicated gRNA‥ **(B)** T7 endonuclease I cleavage assay confirms gRNA-mediated gene editing in transduced cells. Expected digest products are located in Table S4. **(C)** Cell viability assays in various PDAC cell lines stably expressing Cas9, transduced with lentivirus delivering indicated gRNAs. HPAF-II, PaTu8988s and AsPC-1 are sensitive to Wnt pathway inhibition and contain *RNF43* mutations. PANC-1 and BxPC-3 are insensitive to Wnt pathway inhibition. **(D-F)**. RT-qPCR of Wnt target genes (*AXIN2, NKD1*) and differentiation induced gene, *MUC5AC* (n=2), in HPAF-II and PaTu8988s Cas9 cell lines transduced with lentivirus delivering indicated gRNA. All data are represented as means +/- SD, n = 3 independent experiments unless otherwise noted. ***p < 0.001, **p < 0.01, and *p < 0.05, two-tailed unpaired t test, all comparisons to LacZ control gRNA.

Given the potential combinatorial complexity of the WNT pathway (i.e. 19 Wnts and 10 Frizzleds), we were surprised to find that a single Wnt-Fzd ligand-receptor pair is sufficient to drive cellular proliferation in HPAF-II cells. RNA-seq analysis revealed that several of the Wnt genes (*WNT2B, WNT3, WNT7A, WNT7B*, *WNT9A*, *WNT10A*, *WNT10B, WNT16*) and Fzd genes (*FZD1*, *FZD2*, *FZD3*, *FZD4*, *FZD5*, *FZD6*, *FZD7*) are expressed in HPAF-II cells, suggesting that the FZD5-WNT7B circuit is not driven simply by expression (Fig. 3A, Table S5). To further confirm this idea and rule out the possibility of a disconnect between RNA and protein levels for the Wnt receptors in HPAF-II cells, we generated a panel of recombinant antibodies, which we call the ‘FZD profiler’, that can detect and discriminate all but one of the ten Frizzled receptors. Briefly, we used a phage-displayed fragment antigen-binding (Fab) library^22^ and performed binding selections on the purified cysteine rich domains (FZD-CRDs) of each of the 10 human FZD proteins except for FZD3-CRD, which we could not purify (Fig. 3b-d). We chose the most selective Fabs for each of the FZD-CRDs based on Fab-phage ELISAs and converted these to purified Fabs. To characterize the binding specificity of these Fabs on cells, we generated a panel of 10 CHO cell lines each expressing the CRD domain of a different myc-tagged FZD receptor anchored at the plasma membrane through a GPI anchor (CHO-myc-FZD-GPI). Despite the high sequence identity between Frizzled family members (Fig. 3b-c, Fig. S5a-b), we were able to identify selective Fabs for FZD4, FZD5, FZD6 and FZD10 as assessed by immuno-fluorescence and flow cytometry (Fig. 3e and S6). Moreover, we found Fabs that bound to FZD1/7, FZD2/7, FZD5/8, FZD1/2/5/7/8 or FZD4/9/10, which can be used to discriminate expression of the remaining FZDs excluding FZD3 (Fig. 3e and S6). The ‘FZD profiler’ therefore consists of 10 different Fabs that can be used to discriminate expression of 9 different FZDs. The FZD profiler was then used to confirm that HPAF-II cells express FZDs 1,5,6 and possibly 8 (Fig. 3f-g).

**Fig. 3.**
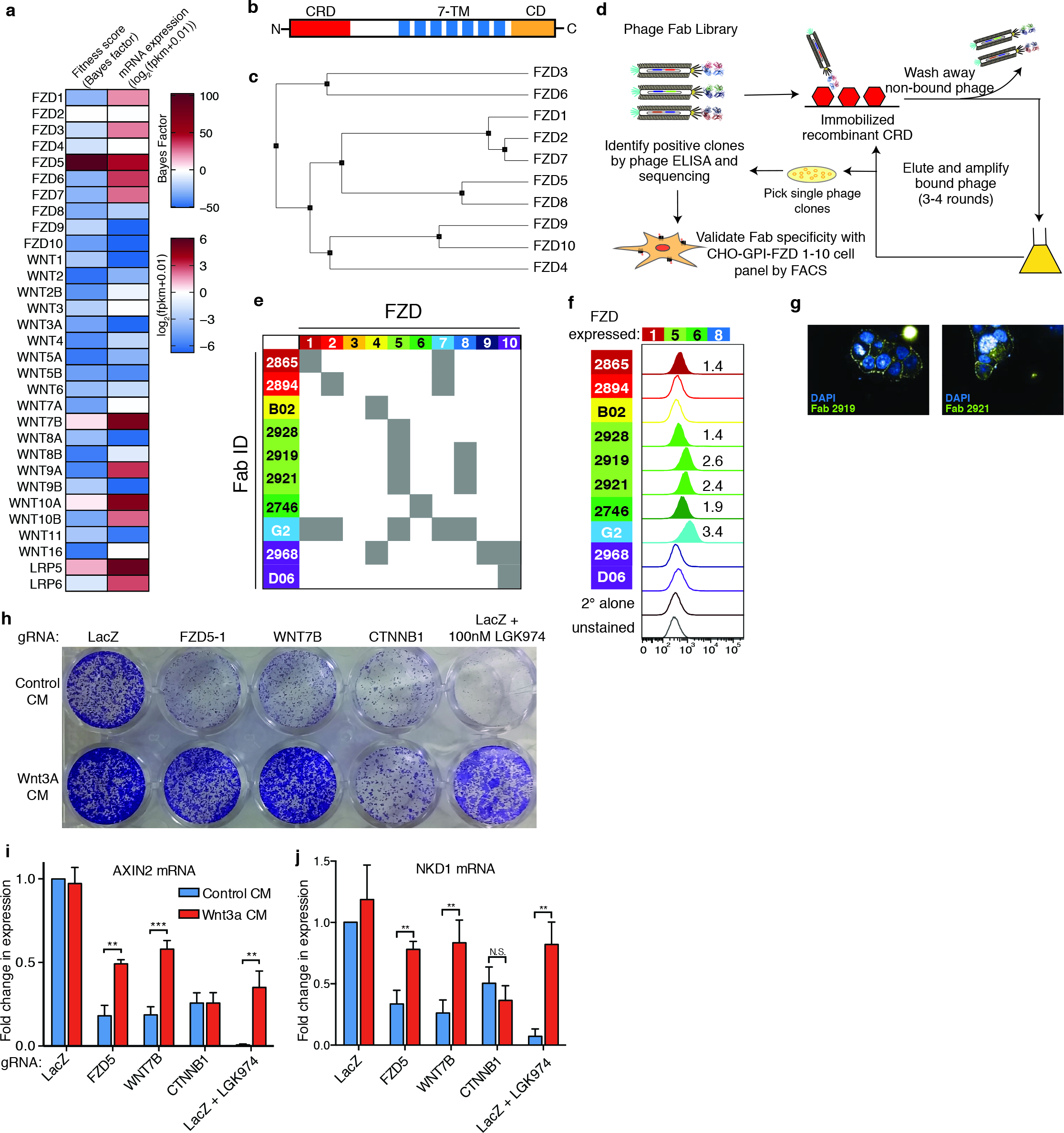
Wnt and Frizzled expression patterns are not predictive of essentiality **(A)** BF and mRNA expression for all *FZD*, *WNT* and *LRP5/6* genes. **(B)** FZD receptors are composed of an extracellular N-terminal cysteine rich domain (CRD), seven-pass transmembrane domain (7TM), and the intracellular C-terminal domain (CD). **(C)** Sequence identity tree among CRD of human FZD receptors (Fig. S5A-B). **(D)** Schematic for Fab selection by phage display. Phage-displayed Fabs are selected for binding to an immobilized FZD CRD of interest. Unbound phages are washed away, and bound phages are amplified over 3-4 rounds to enrich for specific Fabs. Candidate Fabs are validated by flow cytometry using FZD-CRD-GPI overexpressing CHO cell lines for each human FZD. **(E)** Specificity profile of anti-FZD Fabs forming the ‘FZD profiler’. Color of each ID box corresponds to the color of the FZD receptor used to obtain the Fab in selections. **(F)** Determination of FZD receptor membrane expression in HPAF-II cells. Value indicates Median fluorescence intensity (MFI). MFI greater than 1.35X the secondary antibody alone was taken as evidence of endogenous expression. FZD receptors found at the surface of HPAF-II are indicated at the top. **(G)** Indirect immunofluorescence, using Fabs 2919 and 2921 that both detect FZD5 and FZD8, showing cell surface expression in HPAF-II cells. **(H)** Wnt3A conditioned media (CM) rescues proliferation defect and Wnt target gene expression **(I-J)** in *FZD5*, *WNT7B*, but not in *CTNNB1* knockout HPAF-II cells. Mean +/- SD, n = 3 independent experiments. ***p < 0.001, **p < 0.01, two-tailed unpaired t test.

Since multiple Frizzled receptors are expressed on these cells, we predicted that their stimulation with high levels of exogenous Wnt3A would bypass the ligand-receptor pair specificity and this would rescue the growth inhibition seen in *WNT7B* and *FZD5* knockout cells. Confirming this prediction, treatment of *FZD5*or *WNT7B*, but not *CTNNB1* knockout cells with Wnt3A conditioned media (CM) rescued their growth (Fig. 3h), as well as the expression of Wnt target genes *AXIN2* and *NKD1* (Fig. 3i, j). We conclude that FZD5 acts as the receptor for WNT7B to transduce Wnt-β-catenin signaling in *RNF43* mutant PDAC cells.

Based on our results showing that FZD5 was essential for the growth of *RNF43* mutant PDAC cells, we next developed anti-FZD5 full-length human recombinant antibodies (rAbs) and evaluated their binding properties and efficacy in *RNF43* mutant PDAC cells. Using the antibody phage-display system described above for the FZD profiler (Fig. 3d), we isolated a pair of Fabs exhibiting high-affinity binding to human FZD5 CRD that also exhibited cross reactivity to FZD8 CRD, which is the most homologous to FZD5 CRD (Fig. 3c), and converted these Fabs to full-length IgG (IgG-2919 and IgG-2921). Treatment of *RNF43* mutant PDAC cell lines HPAF-II, PaTu8988S and AsPC-1 with the IgGs led to dose dependent growth inhibition (Fig. 4a-c), but they had no effect in PANC-1 and BxPC-3 (Fig. 4d-e). In addition, anti-FZD5 IgG treatment led to inhibition of *AXIN2* and *NKD1* mRNA when compared to control IgG, demonstrating specific Wnt-β-catenin pathway inhibition (Fig. 4f). We also tested the effects of IgG-2919 and IgG-2921 in three patient-derived PDAC cell lines and observed significant anti-proliferative efficacy in GP2A cells, which harbor an *RNF43* variant (R117H) that was previously linked to PDAC in a genome wide association study^23^ and shown to affect RNF43 mRNA instability^24^, but not in GP3A or GP7B (*RNF43* WT) cells (Fig.4g, Fig S7). We next evaluated the efficiency of IgG 2919 to inhibit tumor growth in a subcutaneous xenograft mouse model using HPAF-II cells and showed that twice-weekly dosing at 1 or 2 mg/kg led to 46% or 73% tumor growth inhibition, respectively (Fig. 4h-i, Fig. S8a-b), with no signs of toxicity (Fig. S8c). In addition, histological analysis of the tumors revealed increased mucin production as visualized by Alcian Blue staining, consistent with cellular differentiation (Fig. 4j, S8d)^19^. We conclude that targeting FZD5 with highly specific rAbs represents a new therapeutic opportunity for treatment of *RNF43* mutant cancer.

**Fig. 4.**
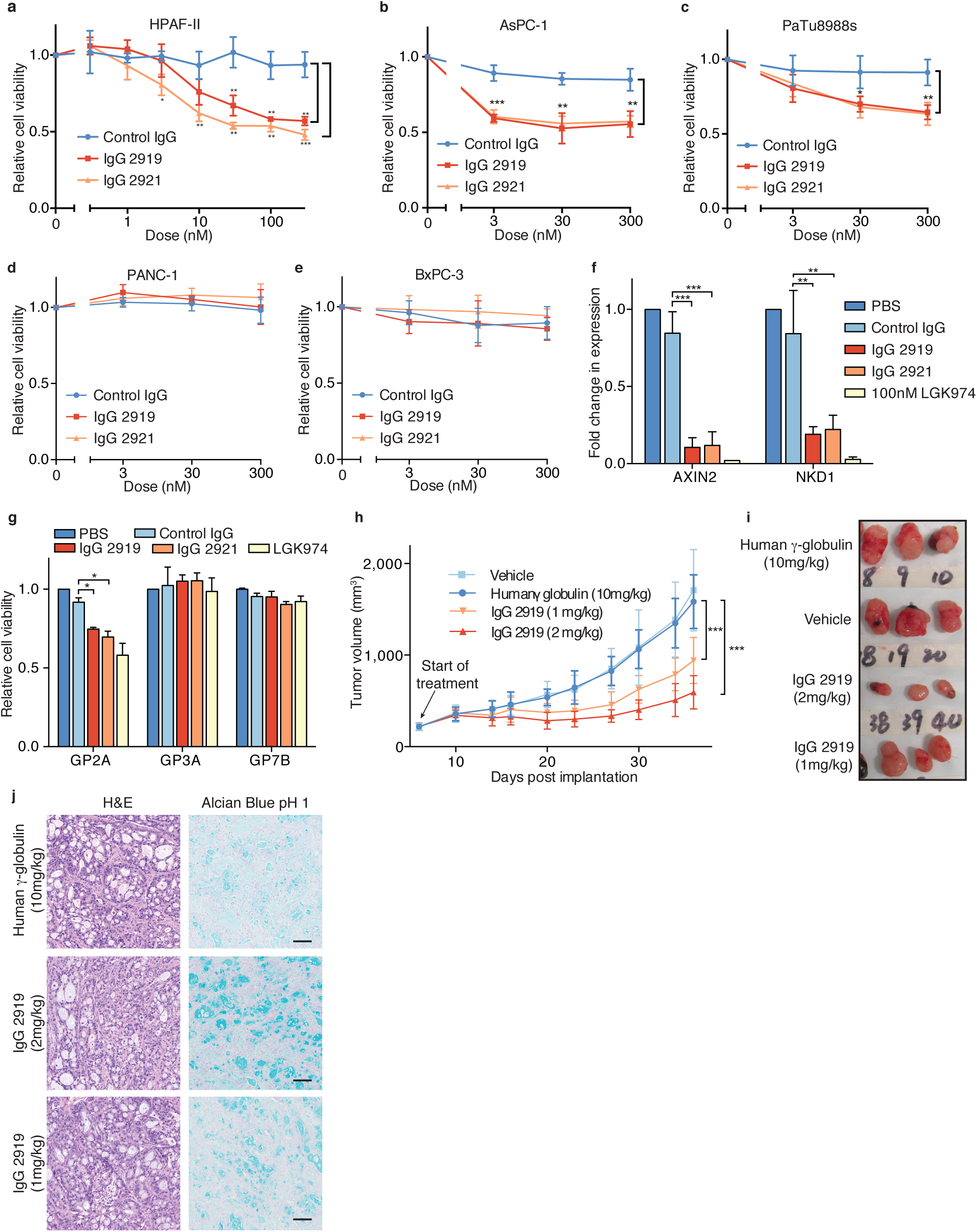
Anti-FZD5 antibodies inhibit growth of *RNF43* mutant PDAC *in vitro* and *in vivo*. **(A-E)** Cell viability assays in *RNF43* mutant and Wnt dependent cell lines (HPAF-II, AsPC-1 and PaTu-8988s and Wnt inhibitor insensitive cell lines (PANC-1 and BxPC-3). Cells were treated with indicated doses of IgG 2919 or 2921, both targeting FZD5. **(F)** RT-qPCR of Wnt-β-catenin target genes *AXIN2* and *NKD1* in HPAF-II after treatment with 300 nM IgG 2919 or 2921. **(G)** Cell viability assays in three patient derived cell lines, each treated with 300nM IgG 2919, 2921 or 100nM LGK974. GP2A contains a *RNF43* variant, whereas GP3A and GP7A are *RNF43* WT. **(H)** *In vivo* treatment with IgG 2919 in HPAF-II mouse intraperitoneal xenografts (n=9 forvehicle, 1mg/kg IgG 2919 conditions, n=10 for human γ-globulin and 2mg/kg IgG 2919 conditions). **(I)** Representative images of xenograft tumors obtained in (H). (**J**) Histological staining of representative tumors from control and antibody-treated xenograft models. All data are represented as means +/- SD, n = 3 independent experiments, unless otherwise noted. ***p < 0.001, **p < 0.01, and *p < 0.05, two-tailed unpaired t test.

## Discussion

Our results revealed a precise requirement for the WNT7B-FZD5 ligand-receptor pair for the growth of PDAC cells harboring *RNF43* mutations. Interestingly, *WNT7B* was previously identified as a physiological regulator of pancreatic progenitor cell growth coordinating pancreas development through autocrine signaling in the epithelium and paracrine response in the surrounding mesenchyme^25^. Gene expression analysis and shRNA-mediated knockdown further supported a role for WNT7B in PDAC cell growth^26^. Our study supports these results and provides evidence that FZD5 acts as the functional receptor for autocrine WNT7B in this context. Whether other stromal Wnt ligands contribute to tumorigenesis in PDAC remains to be determined. Previous attempts at measuring specificity between different Wnt ligands and Frizzled receptors have largely failed, possibly due to overexpression of proteins not being representative of interactions that occur at endogenous doses^27^. Since we showed that several Wnt and Frizzled family members are expressed in these cells, ligand-receptor specificity cannot be explained only by their differential expression and may be governed by small changes in binding affinities that are enough to provide the needed discrimination under normal expression levels. Since stimulation with Wnt3A-CM can rescue the growth of HPAF-II cells and bypass the WNT7B-FZD5 circuit, this indicates that other Frizzled receptors are functional at the surface of the cells and suggests the possibility that upregulation of another Wnt could represent a mechanism of resistance to Frizzled inhibitors.

Despite the known requirement of the Wnt-βcatenin signaling axis in *RNF43* mutated PDAC^19^, the complexity of this pathway in terms of number of Wnt ligands, FZD receptors or intracellular components makes the prediction of therapeutic targets difficult. The identification of specific genetic vulnerabilities using genome-wide CRISPR functional screens therefore provides an unbiased and powerful means to identify context-specific fitness genes that can be harnessed therapeutically for the treatment of human diseases such as cancer. The identification of FZD5 as one of the top essential genes in *RNF43* mutated PDAC and the effectiveness of anti-FZD5 antibodies at inhibiting their growth validated FZD5 as a novel therapeutic target in this context.

By selectively targeting the exact Wnt signaling circuit activated in *RNF43* mutated tumors, FZD specific antibodies may be less toxic than PORCN inhibitors that inhibit all Wnt signaling or promiscuous antibodies that target many Frizzled receptors.

## Methods

### Plasmids

pLCKO lentiviral vector used for TKO library construction and expression of individual gRNAs was constructed as described previously^15^ and deposited with Addgene (#73311). Briefly, sgRNA scaffold was amplified from px330 (Addgene 42230) and cloned into pLKO.TRC005 lentiviral vector (Broad Institute, Cambridge, MA) using AgeI/EcoRI restriction sites. BfuAI sites were added at the 5’ end of the sgRNA scaffold for cloning the TKO library or individual gRNA target sequences. Lenti Cas9-2A-BsdR vector used to generate Cas9 stable expression cell lines was constructed as described previously^15^ (Addgene 73310). Briefly, lentiCRISPR pXPR_001 (Addgene 49535) was modified through removal of the sgRNA scaffold region through NdeI/EcoRI digest and blunt ends generation using Large Klenow Fragment (NEB). 2A-puro was replaced with 2A-BsdR by PCR using the NheI/MluI sites. Each of the 10 human FZD CRDs were cloned into the lentiviral expression plasmid pLenti-puro in the following cassette: FZD_CRD-MYC-GPI.

### Cell Culture

HPAF-II, PaTU-8988s, PANC-1, HEK293T and mouse L-cell cell lines were maintained in DMEM conatining 4.5g/L D-glucose, L-glutamine (ThermoFisher #11965) and supplemented with 10% FBS (ThermoFisher) and Penicillin/Streptomycin (ThermoFisher #15140-163). AsPC-1 and BxPC-3 cell lines were maintained in RPMI 1640 with L-glutamine (ThermoFisher #11875) supplemented with 10% FBS and Penicillin/Streptomycin. CHO cells were maintained in DMEM/F12 (ThermoFisher #11320-033), supplemented with 10%FBS and penicillin/streptomycin. All cell lines were maintained at 37° and 5% CO_2_. For puromycin selection cells were selected in medium containing 2μg/ml puromycin dihydrochloride (Bioshop Canada #PUR333). For blasticidin selection cells were selected in medium containing 10μg/ml (HPAFII), 8μg/ml (PaTu-8988s), 5μg/ml (PANC-1), 2μg/ml (BxPC-3) or 1μg/ml (AsPC-1) of blasticidin hydrochloride (Bioshop Canada #BLA477). Cells were tested for mycoplasma with MycoAlert mycoplasma detection kit (Lonza # LT07-118) before use in experiments.

### gRNA Library Design and Construction

gRNA library design and construction were described previously^15^. Briefly, ~90k gRNA candidate sequences were chosen based on minimal off-target sites and optimized cleavage efficiency. The library was designed to target as many protein-coding exons as possible, with a maximum of 6 gRNA/gene. This yielded a library targeting 17232 genes. The library was synthesized in a pooled oligo array of 58-mers (CustomArray), with each gRNA target flanked by BfuAI restriction sites (oligo sequence below). The oligo pool was PCR amplified (primers listed below), purified (Qiaquick nucleotide removal kit, Qiagen #28304) and ligated into pLCKO in a one-step digestion/ligation reaction with BfuAI and T4 ligase (NEB). Ligation products were purified with Qiaquick nucleotide removal kit (Qiagen #28304), transformed in Electromax Stbl4 competent cells (ThermoFisher #11635-018) and grown on LB-Carbenicillin (100μg/ml, ThermoFisher #10177-012) plates. >5.8E7 colonies were harvested, for ~650-fold library coverage. Plasmid DNA was extracted from colonies with QIAfilter Plasmid Giga Kit (Qiagen #12291).

> *Oligo array template:*
>
> AGAGAACCTGCAGAGACCGNNNNNNNNNNNNNNNNNNNNGTTTAGAGGCAGGTAGAGA
>
> *Primers for amplifying CRISPR library:*
>
> Forward: TGTCAGTTGTCATTCGCGAAAAAGAGAACCTGCAGAGACC
>
> Reverse: GTCACTGACGCGGTTTTGTAGATCTCTACCTGCCTCTAAA

### Lentiviral Production

Lentivirus production of the gRNA library was completed as described previously^15^. Briefly, 9 million HEK293T cells were seeded per 15cm plate. 24-hours post seeding, cells were transfected with a mixture of 14μg pLCKO gRNA library plasmid pool, 16μg packaging vector psPAX2 (Addgene 12260), 1.56μg envelope vector pMD2.G (Addgene 12259), 93.6μl X-treme Gene transfection reagent (Roche #06366236001) and 1.4ml Opti-MEM medium (ThermoFisher #31985-070), per plate. 24 hours post transfection medium was replaced with DMEM, 1.1g/100ml BSA, Penicillin/Streptomycin. Viral media was harvested at 48 and 72 hours post transfection by centrifugation at 1500RPM for 5 minutes at 4°, aliquoted and and filtered before freezing at -−80°.

For routine lentiviral production, 3.5 million HEK293T cells were seeded in 10cm plates 24 hours before transfection. 5μg of lentiviral delivery vector (pLCKO or LentiCas9), 4.5μg psPAX2 and 0.5μg pMD2.G were transfected per plate with Lipofectamine 2000 (ThermoFisher #11668-019), following manufacturers protocol. 24 hours post transfection medium was changed. Viral media was harvested 48 hours post transfection by centrifugation at 2000rcf for 5 minutes followed by filtering through a 0.2μm syringe filter.

### Lentiviral Transduction and MOI Determination

Cells were seeded at low density (for three days growth) in medium containing between 0.125% and 8% viral medium and 8μg/ml polybrene (Sigma #H9268-5G). 24 hours post transduction, cells were placed in selective medium containing puromycin. Multiplicity of infection (MOI) determination was done by comparing cell counts in control and puromycin containing wells, transduced at various viral medium concentrations after 48 hours puromycin selection.

### Lentiviral gRNA Library Essentiality Screen in HPAF-II

HPAF-II cell line was transduced with Lenti Cas9-2A-BsdR as described above and selected in 10μg/ml blasticidin. A polyclonal stable cell line was established and single clones were isolated by limited dilution. Clones were expanded and screened for Cas9 expression via western blot and cleavage activity with pLCKO delivered gRNAs (data not shown).

The selected HPAF-II Cas9-2A-BsdR clone was transduced with the 90k gRNA library at an MOI of 0.3 and a library fold-coverage of 300x (~27 million transduced cells). 72 hours post-infection (and 48 hours post puromycin selection) cells were split into two independent replicate populations of minimum 200-fold library coverage (18 million cells). In addition, T0 reference samples were collected (18 million cells) for genomic DNA extraction. Replicate populations were passaged in parallel every four days, with 18 million cells seeded over five 15cm plates per population. Samples were collected at T15, T27, T31 and T35, at approximately 10, 18, 21 and 23 doublings respectively.

### Screen sample preparation for sequencing

Genomic DNA was extracted and prepared for PCR as described previously^28^. Briefly, genomic DNA was extracted with QiaAmp DNA Maxi Kit (Qiagen #51192), following manufacturers protocol. Following genomic DNA extraction, the DNA was ethanol precipitated and resuspended in 10mM Tris-HCl pH8.5 at a concentration greater than 500ng/μl.

2-steps nested PCR amplification of gRNA target sequences for Illumina sequencing was completed as described previously^15^. Briefly, 50μg of genomic DNA per sample was used as template for amplification using primers listed below. This was completed with KAPA HiFi polymerase (Kapa Biosystems #KK2602) and split over ten 50μl reactions. After amplification, reactions were pooled and 5μl was used as template for amplification with primers containing Illumina TruSeq adapters. Final PCR products were gel purified with Purelink combo kit (ThermoFisher #K2200-01). Sequencing was completed with Illumina HiSeq2500, as described previously^15^.

> *Step 1 PCR Forward:* AGGGCCTATTTCCCATGATTCCTT
>
> *Step 1 PCR Reverse*: TCAAAAAAGCACCGACTCGG
>
> *TruSeq adapters with i5 barcodes:*
>
> AATGATACGGCGACCACCGAGATCTACACTATAGCCTACACTCTTTCCCTACACGACGCTCTTCC GATCTTGTGGAAGGACGAGGTACCG
>
> AATGATACGGCGACCACCGAGATCTACACATAGAGGCACACTCTTTCCCTACACGACGCTCTTCC GATCTTGTGGAAGGACGAGGTACCG
>
> AATGATACGGCGACCACCGAGATCTACACCCTATCCTACACTCTTTCCCTACACGACGCTCTTCCG ATCTTGTGGAAGGACGAGGTACCG
>
> AATGATACGGCGACCACCGAGATCTACACGGCTCTGAACACTCTTTCCCTACACGACGCTCTTCC GATCTTGTGGAAGGACGAGGTACCG
>
> *TruSeq adapters with i7 barcodes:*
>
> CAAGCAGAAGACGGCATACGAGATCGAGTAATGTGACTGGAGTTCAGACGTGTGCTCTTCCGATC TATTTTAACTTGCTATTTCTAGCTCTAAAAC
>
> CAAGCAGAAGACGGCATACGAGATTCTCCGGAGTGACTGGAGTTCAGACGTGTGCTCTTCCGATC TATTTTAACTTGCTATTTCTAGCTCTAAAAC
>
> CAAGCAGAAGACGGCATACGAGATAATGAGCGGTGACTGGAGTTCAGACGTGTGCTCTTCCGAT CTATTTTAACTTGCTATTTCTAGCTCTAAAAC
>
> CAAGCAGAAGACGGCATACGAGATGGAATCTCGTGACTGGAGTTCAGACGTGTGCTCTTCCGATC TATTTTAACTTGCTATTTCTAGCTCTAAAAC

### Analysis of CRISPR Negative Selection Screen

Read counts for each gRNA were normalized for each replicate at each of the indicated time points (T27, T31, T35) and a log fold change relative to control (T0) was calculated. The BAGEL algorithm^21^ was used to calculate a Bayes Factor for each gene, representing a confidence measure that the gene knockout results in a fitness defect. Bayes Factors at each time points were summed to a final BF for each gene.

### Gene ontology enrichment analysis

Gene ontology enrichment was completed using GOrilla^29^, using differential Z-score (Fig. 1B, Table S3) for ranking. Results were filtered for FDR<0.05, number of genes used in enrichment ≥5 and enrichment score >3.

### Cloning of individual gRNA target sequences into pLCKO

pLCKO vector was digested with BfuAI (NEB) and gel purified with Purelink combo kit (ThermoFisher #K2200-01). Forward and reverse oligonucleotides coding for the gRNA targets (listed below) were phosphorylated and annealed. Oligos were first phosphorylated with PNK (ThermoFisher #AM2310) and annealed through 95° incubation for 10 minutes followed by slope ramp-down to room temperature. Phosphorylated/annealed oligo pairs were ligated into BfuAI digested pLCKO in a 1:5 molar ratio with T4 ligase (NEB #M0202L) and transformed in DH5α cells. DNA was prepped with GeneElute HP plasmid midi-prep kit (Sigma #NA0200) and verified by Sanger sequencing. Note that gRNA targets were chosen from the 90k TKO library if they were shown to be functional in the screen (FZD5, WNT7B) or through CRISPR design tool (http://crispr.mit.edu/)^30^ (e.g. FZD7, FZD4, FZD8).

> *All gRNA target oligos were designed as follows:*
>
> 5’ – ACCGNNNNNNNNNNNNNNNNNNNN -−3’
>
> 3’- NNNNNNNNNNNNNNNNNNNNCAAA –5’
>
> *Target Sequences:*
>
> LacZ: CCCGAATCTCTATCGTGCGG
>
> CTNNB1: GAAAAGCGGCTGTTAGTCAC
>
> FZD4-1: AGCTCGTGCCCAACCAGGTT
>
> FZD4-2: ATGCCGCCGCATGGGCCAAT
>
> FZD5-1: AGGCCACCACAATGCTGGCG
>
> FZD5-2: TCCGCACCTTGTTGTAGAGC
>
> FZD7-1: GCCGGGGCGCAGCCGTACCA
>
> FZD7-2: TGGTACGGCTGCGCCCCGGC
>
> FZD8-1: TAGCCGATGCCCTTACACAG
>
> FZD8-2: CAACCACGACACGCAAGACG
>
> WNT7B: GGCTGCGACCGCGAGAAGCA

### T7 endonuclease I assay to assess Cas9-gRNA cleavage

5-7 days post pLCKO lentiviral transduction genomic DNA was extracted using PureLink genomic DNA mini kit (ThermoFisher ThermoFisher #K2200-01). Genomic DNA was used as template to amplify targeted locus (primer pairs listed below) using Kapa HiFi polymerase (Kapa Biosystems #KK2602), following manufacturer’s protocol. PCR products were purified with PureLink combo kit (ThermoFisher #K2200-01). DNA concentration in purified PCR products were quantified with Nanodrop 1000 (Thermo Scientific). 200ng of CRISPR edited PCR product was mixed with 200ng of wild-type PCR product with 1x NEB buffer 2.0 for a final volume of 19.5μl. Samples were heated to 95° for 10 minutes, followed by slow ramp-down to room temperature for heteroduplex formation. 0.5μl of T7 endonuclease I (NEB #M0302L) was added to each sample and incubated at 37° for 20 minutes. Immediately following digest, samples were resolved on a 2% agarose gel (BioShop Canada #AGA001.500) containing ethidium bromide (BioShop Canada #ETB444.1). All images are representative of 2 independent experiments.

> FZD4 exon1 forward: TGTCTCCTTCGGGCTAGGAT
>
> FZD4 exon1 reverse: CGGGACGTCTAAAATCCCACA
>
> FZD4 exon2 forward: CAGGTTCTGCTGCCTCTTCA
>
> FZD4 exon2 reverse: AGTGTTGTGCAAAGAGGGCT
>
> FZD5 forward: TTGCCCGACCAGATCCAGAC
>
> FZD5 reverse: TCTGTCTGCCCGACTACCAC
>
> FZD7 forward: TGAGGACTCTCATGCGTCGG
>
> FZD7 reverse: AGCCGTCCGACGTGTTCT
>
> FZD8 forward: TGTCTTGACGCGGTTGTAGAG
>
> FZD8 reverse: TAGATTATCGGCAGACCCCC

### Crystal Violet Staining Proliferation Assay

HPAFII Cas9 cells were transduced with lentivirus generated with the indicated pLCKO plasmid as described above. 24 hours after infection cells were treated with puromycin. After 48 hours of selection, cells were PBS washed extensively, dissociated and counted. 2000 cells per well were re-seeded in 24-well format in media without puromycin. 24 hours post seeding, indicated wells were treated with DMSO control or 100nM LGK-974 (Cayman Chemical #14072) (note that these wells were from the LacZ gRNA population). Medium was renewed every 3-4 days and cells were fixed, 10 days post plating, using 100% ice-cold methanol. After fixation cells were stained with 0.5% crystal violet, 25% methanol solution for 20 minutes at room temperature, after which staining solution was removed and plates were washed several times in dH_2_O. For Wnt3A conditioned media rescue experiment, cells were treated with 25% control or Wnt3A CM from initial infection onwards.

### Cell Viability Assays

For FZD5/7 gRNA experiments, Cas9 expressing stable cell lines were transduced with indicated lentivirus as described above. 24 hours after infection cells were treated with puromycin. After 48-72 hours of puromycin selection, wells were washed with PBS extensively, dissociated and counted. Cells were re-seeded at 1000 cells per well, six wells per gRNA, in 96 well plates. Medium was changed every 3-4 days and viability was measured with Alamar Blue (ThermoFisher #DAL1025) 7-11 days post plating. Briefly, 10ul of Alamar Blue was added to 100μl medium per well and incubated 3-4 hours at 37°, 5% CO_2_. Fluorescence was measured at 560nm excitation, 590nm emission with Spectramax Gemini XS plate reader (Molecular Devices).

For antibody treatments, cells were seeded at 1000-2000 cells per well in 96-well plates. 24 hours after seeding, cells were treated with antibodies in quadruplicates, at the indicated concentrations. Medium was changed and antibodies renewed after 3 days. Viability was measured with Alamar Blue, 6 days after plating, using the same procedure described above.

### Reverse transcription and quantitative real-time PCR

After indicated treatments, cells were lysed in Tri-reagent (BioShop Canada #TSS120) and RNA extracted using the manufacturer’s protocol. RNA concentration was quantified with Nanodrop1000 (Thermo Scientific) and 2μg of RNA per sample was DNase I treated (ThermoFisher #AM2222). DNase treated RNA was used to make cDNA with High-Capacity cDNA Reverse Transcription Kit (ThermoFisher #4368813). Real-time PCR was performed using Power SYBR Green Master Mix on the 7900HT Fast Real-Time PCR system. Primer pairs are listed below. Analysis was done using the comparative cycle threshold (*CT*) method^31^ with all samples normalized to *PPIB* (cyclophilin B) expression.

> PPIB Forward: GGAGATGGCACAGGAGGAA
>
> PPIB Reverse: GCCCGTAGTGCTTCAGTTT
>
> AXIN2 Forward: CTCCCCACCTTGAATGAAGA
>
> AXIN2 Reverse: TGGCTGGTGCAAAGACATAG
>
> NKD1 Forward: TGAGAAGATGGAGAGAGTGAGCGA
>
> NKD1 Reverse: GGTGACCTTGCCGTTGTTGTCAAA
>
> MUC5AC Forward: AGCCGGGAACCTACTACTCG
>
> MUC5AC Reverse: AAGTGGTCATAGGCTTCGTGC

### RNAseq

RNAseq for the HPAF-II cell line was completed as described in detail previously for other cell lines^15^. Briefly, total RNA was extracted using Tri-reagent (BioShop Canada #TSS120) following manufacturers instructions. Sequencing libraries were prepared with Illumina TruSeq V2 RNA library preparation kit. Libraries were sequenced in single reads, 61bp, on a High Output Illumina NextSeq500 flowcell (version1 chemistry). Reads were mapped using Gencode v19 gene models in Tophat v2.0.4. Gene expression values were determined using Cufflinks v2.2.1.

### Isolation and characterization of Fabs against FZD

The anti-FZD Fabs were isolated from a synthetic human Fab phage-displayed library (Library F)^22^. Binding selections, phage ELISAs and Fab protein purification were performed as described^32-34^. Briefly, phage particles displaying the Fabs from Library F were cycled through rounds of panning with purified FZD-Fc fusion (R&D Systems) immobilized on 96-well Maxisorp Immunoplates (Fisher Scientific, Nepean, ON, Canada) as the capture target. After four rounds of selection, phage were produced from individual clones grown in a 96-well format and phage ELISAs were performed to detect specific binding clones. Clones with positive binding were subjected to DNA sequencing. The DNAs encoding for variable heavy-and light-chain domains of the positive binders were cloned into vectors designed for production of Fabs or light chain or IgG1 heavy chain, respectively. and Fabs were expressed from bacterial cells and IgGs from 293F cells (Invivogen, San Diego, CA, USA). Fab and IgG proteins were affinity-purified on Protein A affinity columns (GE Healthcare, Mississauga, ON, Canada).

### Flow cytometry

Primary staining of cells was performed with a 200nM FZD profiler Fab. Alexa Fluor 488 AffiniPure F(ab’)_2_ was used as the secondary antibody (Jackson ImmunoResearch # 109-546-097). c-Myc (9E10) IgG1 (primary antibody, Santa Cruz, lot # D0306) and Alexa Fluor 488 IgG (secondary antibody, Life technologies, lot #1458649) were used as controls. Dead cells were excluded by staining with Fixable Viability Dye eFluor 660 (eBioscience, catalogue number 65-0864). All reagents were used as per manufacturer’s instructions. Flow cytometry was performed on a BD FACSCanto II flow cytometer (BD Biosciences), and data were analyzed with FlowJo software (FLowJo, LLC).

### Wnt3a and control conditioned media

Mouse fibroblast L cells (ATCC #CRL-2648) and human Wnt3A expressing L cells (ATCC #CRL-2647) were grown to near confluence on 15-cm plates. They were then split 1:12 into 25ml of complete medium per plate (15-cm). Medium was collected and 0.2μm filtered 4 and 7 days post seeding.

### Mouse xenograft studies

CB-17 Fox Chase SCID mice (6 weeks old, female) were purchased from Charles River Laboratories (St. Constant, QC, Canada). The mice were housed in a pathogen-free environment at the animal facility at the University of Toronto. The study was conducted according to the guidelines of the Canadian Council on Animal Care (CCAC) and the animal use protocols approved by the University Animal Care Committee (UACC) at the University of Toronto.

The recombinant antibody, IgG 2919, was developed and purified as described above. Human γ globulin was purchased from Jackson ImmunoResearch Laboratories, Inc. (West Grove, PA, USA), and Dulbecco’s phosphate-buffered saline (DPBS, no calcium, no magnesium) was obtained from Thermo Fisher Scientific Inc. (Burlington, ON, Canada). Human γ globulin and D-PBS were used as the experimental controls in this study.

HPAF-II cells were inoculated subcutaneously into the flank of the CB-17 SCID mice with 3 x 10^6^ cells in D-PBS per mouse. Tumor volumes were measured using vernier calipers and the mice were weighed twice weekly. Tumor volume was calculated using the formula: ½ (Length × Width^2^). When tumors reached approximately 200 mm^3^, the mice were randomized into four groups of nine or ten mice each. Each group received one of the following treatments: Human γ globulin (10 mg/kg), D-PBS (15 mL/kg), IgG 2919 (2 mg/kg), or IgG 2919 (1 mg/kg), twice weekly via intraperitoneal injection for four and a half weeks.

For calculation of percentage of tumor growth inhibition (TGI), groups treated with antibody (Ab test) were compared with group treated with human γ globulin (control). TGI (%) was calculated using the formula: TGI (%) = {(mean TVG control – mean TVG Ab test) /mean TVG control} x 100, where the mean TVG (tumor volume growth) = mean tumor volume at a defined study day – mean tumor volume the day of the first dosing. Statistical significance was examined by Student’s t-test (two-tailed). P-values less than 0.05 were considered statistically significant.

### Histological staining

Tumor staining was carried out at the immuno-histopathology and tissue processing lab at the University Health Network. Briefly, three representative tumors from each treatment group (human γ-globulin at 10mg/kg, IgG-2919 at 1mg/kg and IgG-2919 at 2mg/kg) were embedded into a wax block and paraffin embedded tumors were cut into thin sections and mounted onto a microscope slide for routine staining with Hematoxylin and Eosin, Periodic Acid Schiff or PAS (Abcam – ab150680), Alcian Blue pH 1.0 (Abcam – ab150661), and Alcian Blue pH 2.5 (Abcam – ab150662). An Axioscan slide scanner system was used to generate high resolution digital images of the whole tumor sections at 40x in brightfield mode, and the images were exported as.png files using ZEN software.

#### Sanger sequencing

RNF43 exon 2 locus was amplified by PCR from GP2A genomic DNA using primer pairs listed below. Sequencing was completed at The Centre for Applied Genomics (Toronto, ON) with a primer listed below.

> RNF43 exon 2 forward: TGGTGCTAGTCGGAGGAGAA
>
> RNF43 exon 2 reverse: GTTGCTTTGCAGCAAGCTCT
>
> Sequencing primer: CCTGCCTGGTACCTCCCTAG

#### Statistical analysis

All statistical analyses were completed with Graphpad Prism 6 (GraphPad Software, La Jolla California USA).

## Acknowledgments

We wish to thank members of the Angers and Moffat labs for discussions. We are thankful to Dr. David Hedley for providing the patient-derived xenograft cells. This work was supported from grants funded by the Canadian Institutes for Health Research to S.A. (CIHR-273548) and J.M. (CIHR-342551) and the Ontario Research Fund to S.S‥ J.M. holds a Canada Research Chair in Functional Genomics of Cancer and S.A. holds a Canada Research Chair in Functional Architecture of Signal Transduction.

## Author Contributions

Conceptualization, S.A., J.M., S.S.; Investigation, Z.S., T.H., M.C., Z.P., M.R., X.W., J.A., and J.P.; Writing, S.A., Z.S., J.M.; Funding Acquisition, S.A., J.M., S.S; Supervision, S.A., J.M., S.S, and J.P.

